# Non3nExonFinder: transcript-aware identification of shared non-triplet coding exons

**DOI:** 10.64898/2026.09.22.753378

**Authors:** Sung-Yeon Lee, Joo-Hee Lee

## Abstract

Selection of an appropriate coding exon is an important upstream step in exon deletion-based loss-of-function strategies, particularly for genes with multiple transcript isoforms. However, manual comparison of exon structures across multiple transcripts can be time-consuming and may overlook transcript-specific differences in exon boundaries and coding sequence (CDS) composition. Non3nExonFinder was developed as an interactive R/Shiny-based tool for identifying shared non-triplet coding exons across user-defined human or mouse NCBI RefSeq protein-coding transcripts. Non3nExonFinder evaluates exon/CDS annotations and coding contributions for each transcript, identifies common exons with identical genomic boundaries, and prioritizes candidates based on CDS-only status and coding-length divisibility by three. Non-triplet CDS-only exons that are internal in all selected transcripts are classified as high-confidence candidates. Independent reconstruction from the original RefSeq GTF files across 80 genes (40 human and 40 mouse), comprising 6,848 transcript-specific exon instances, showed complete concordance with Non3nExonFinder outputs, with exact recovery of all candidate sets. Non3nExonFinder provides a reproducible upstream framework for transcript-aware prioritization of exon-deletion targets.

## INTRODUCTION

Targeted deletion of coding exons is widely used in loss-of-function studies, including conditional knockout strategies (Bradley et al. 2012; Skarnes et al. 2011). In CRISPR/Cas9-based approaches, single-guide RNAs (sgRNAs) can be positioned in intronic regions flanking the target exon to remove the intervening genomic segment (Bonafont et al. 2019; Lee et al. 2026; Peterson et al. 2017). When the number of coding nucleotides removed is not divisible by three, exon deletion can induce a frameshift in the downstream coding sequence (CDS), potentially leading to the formation of a premature termination codon (Peterson et al. 2017; Skarnes et al. 2011). Accordingly, selection of an appropriate non-triplet coding exon can provide an important starting point for designing exon-deletion experiments.

In practice, application of this criterion is more complex for genes with multiple transcript isoforms, such as *DMD, TTN*, and *CACNA1C*, because target exon selection requires consideration of isoform-specific exon usage (Jiang et al. 2024; Roberts et al. 2015; Tuffery-Giraud et al. 2017). For example, an exon included in the CDS of one transcript may be absent from another transcript, and even the same genomic exon may contain untranslated sequence or contribute a different number of coding nucleotides depending on transcript-specific CDS boundaries (Frankish et al. 2021). Thus, selecting a target exon based on a single representative transcript may fail to account for alternative RefSeq isoforms with distinct coding potential (Morales et al. 2022). Given these complexities, manual comparison of exon structures across multiple transcripts is feasible for relatively simple genes, but becomes increasingly laborious and error-prone as the number of annotated transcripts increases.

To support such design tasks, several computational tools have been developed for CRISPR guide design and, in some cases, exon-deletion strategy design (Kuno et al. 2019; Labun et al. 2019; Malekos et al. 2025; Peterson et al. 2017). These tools provide useful functions such as guide-RNA design, identification of regions shared among multiple isoforms, and prediction of reading-frame consequences. However, these existing tools do not simultaneously integrate user-defined RefSeq transcript selection, exact shared-exon boundary detection, transcript-specific CDS contribution analysis, and assessment of coding-length divisibility across all selected transcripts.

This study presents Non3nExonFinder, an interactive tool for identifying shared non-triplet coding exons across user-selected human or mouse RefSeq transcripts. The tool uses version-specific RefSeq-derived annotations to compare exon and CDS information across individual transcripts and identifies common exons with identical genomic boundaries among all selected transcripts. Common exons are subsequently evaluated according to CDS composition and coding-length divisibility by three, with internal CDS-only non-triplet exons reported as high-confidence candidates. Non3nExonFinder therefore provides a reproducible upstream annotation framework for prioritizing candidate exon-deletion targets before downstream CRISPR guide design.

## METHODS

### Software implementation and annotation resources

Non3nExonFinder was implemented in R v4.3.3, with the user interface developed using Shiny v1.12.0 (Chang et al. 2025). Transcript, exon, and CDS annotations were accessed using the GenomicFeatures and AnnotationDbi packages (Huber et al. 2015; Lawrence et al. 2013), with dplyr and DT used for data processing and interactive table display, respectively. The analysis was restricted to human and mouse NCBI RefSeq protein-coding transcripts represented by NM_ accessions (O’Leary et al. 2016). RefSeq-derived TxDb databases based on GRCh38.p14 for human and GRCm39 for mouse were stored locally, thereby providing a fixed annotation environment without requiring live annotation queries during routine analysis.

### Transcript-specific exon annotation and candidate identification

For each analysis, exon and CDS annotations were queried only for the selected transcripts rather than loading transcript structures from the entire TxDb database. For each exon, chromosome, genomic start and end coordinates, strand, and exon rank were retrieved. CDS intervals were restricted to those belonging to the same transcript, chromosome, and strand and overlapping the genomic coordinates of the corresponding exon. The transcript-specific coding contribution of each exon (cds_bp) was calculated by summing the lengths of the genomic overlaps between the exon and annotated CDS intervals.

Genomic exon length was calculated from the start and end coordinates, and each exon was classified as untranslated region (UTR)-only, CDS-only, or mixed according to its coding contribution. Exons with no CDS overlap were classified as UTR-only, those entirely covered by CDS were classified as CDS-only, and those containing both coding and noncoding sequence were classified as mixed. For exons containing CDS, coding-length divisibility was evaluated as cds_bp mod 3, and an exon was classified as non-triplet when cds_bp was greater than zero and not divisible by three. An exon was classified as internal when its exon rank was greater than 1 and lower than the maximum exon rank of the corresponding transcript.

A common exon was defined as an exon with identical chromosome, genomic start coordinate, genomic end coordinate, and strand across all selected transcripts. Common exons were then classified sequentially into four categories: (i) all common exons, (ii) common exons classified as CDS-only in every transcript, (iii) common CDS-only exons with coding contributions not divisible by three in every transcript, and (iv) high-confidence candidates that additionally represented internal exons in every transcript.

### User interface and output

The Shiny interface allows users to select either human or mouse annotation and enter one or more RefSeq transcript accessions. For an entered transcript, users can additionally retrieve other eligible RefSeq protein-coding transcripts associated with the same Gene ID. Results are displayed in separate interactive tables for high-confidence candidates, common non-triplet CDS-only exons, common CDS-only exons, all common exons, and transcript-specific exon annotations. The output includes transcript-specific exon rank, coding contribution, CDS/UTR classification, and candidate information, and the result tables can be exported in comma-separated values (CSV) format. The implementation queries only the selected transcripts and independently calculates CDS overlap and modulo-three status for each transcript.

### Representative *TIA1* analysis and independent validation

As a representative multi-transcript use case, eight human RefSeq protein-coding transcripts of *TIA1*(NM_001351508.2, NM_001351509.2, NM_001351510.2, NM_001351511.1, NM_001351512.1,NM_001351513.1, NM_001351514.2, and NM_001351515.2) were analyzed together. Transcript-level annotations and candidate sets generated by Non3nExonFinder were compared with results independently reconstructed from the original NCBI RefSeq Gene Transfer Format (GTF) file (GCF_000001405.40_GRCh38.p14_genomic.gtf.gz).

For independent validation, exon structure and coding contribution were reconstructed directly from the exon, CDS, and stop_codon features of the original GTF without using the TxDb object or the annotation functions implemented in Non3nExonFinder. Stop-codon features were included together with CDS features to ensure consistency with the coding-interval representation used by the TxDb annotation. The reconstructed data were used to independently derive exon coordinates and rank, coding contribution, CDS/UTR classification, modulo-three status, non-triplet status, and internal-exon status. Candidate sets were then reconstructed using the same four sequential filtering criteria and compared directly with the corresponding Non3nExonFinder outputs.

### Large-scale independent validation

To evaluate the performance of Non3nExonFinder across diverse transcript structures, 40 multi-transcript genes were randomly selected from each of the human and mouse annotations. Eligible genes were required to contain at least two RefSeq transcript accessions matching the pattern ^NM_[0-9]+(\.[0-9]+)?$, and all eligible transcripts from each selected gene were included in the analysis. *TIA1* was excluded from the human sampling pool because it was evaluated separately as the representative example. Fixed random seeds of 260921 and 260922 were used for human and mouse sampling, respectively.

Independent validation was performed using the original NCBI RefSeq GTF files for human GRCh38.p14 (GCF_000001405.40_GRCh38.p14_genomic.gtf.gz) and mouse GRCm39 (GCF_000001635.27_GRCm39_genomic.gtf.gz). The exon, CDS, and stop_codon features corresponding to the selected transcripts were extracted directly from the original GTF files, and exon rank was reconstructed from the exon_number attribute. Coding contribution was independently calculated from the genomic overlap between each exon and the union of CDS and stop-codon intervals belonging to the same transcript, chromosome, and strand. CDS/UTR classification, modulo-three status, non-triplet status, and internal-exon status were subsequently derived without using the classification functions implemented in Non3nExonFinder.

For exon-level validation, the independently reconstructed raw-GTF annotations were matched to the corresponding Non3nExonFinder outputs by transcript and exon rank. Concordance was assessed for exon coordinates and rank, coding contribution, CDS/UTR classification, coding-length modulo-three status, non-triplet classification, and internal-exon status. For candidate-level validation, the sets of all common exons, common CDS-only exons, common non-triplet exons, and high-confidence candidates were independently reconstructed from the raw GTF for each gene and compared with the corresponding Non3nExonFinder candidate sets for exact agreement.

### Computational performance assessment

Computational performance was evaluated using a human gene containing at least 40 eligible NM_ transcripts. Among genes meeting this criterion, the gene with a transcript count closest to 40 was selected deterministically, and nested transcript sets containing 1, 5, 10, 20, and 40 transcripts were generated from the consistently ordered transcript list. For each transcript-set size, one untimed warm-up analysis was performed, followed by 10 timed runs. Elapsed execution time was measured using the R system.time() function and summarized as the median and interquartile range for each transcript-set size.

## RESULTS

### Non3nExonFinder provides a transcript-aware workflow for exon prioritization

Non3nExonFinder provides an interactive workflow for transcript-aware identification of shared coding exons across user-selected human or mouse RefSeq transcripts. The overall workflow comprises transcript input and annotation retrieval, transcript-specific exon/CDS analysis, sequential candidate filtering, and interactive output (**Fig. 1a–d**). A comparison of the principal features of Non3nExonFinder with selected existing CRISPR-related tools is summarized in **Table 1**.

**Table 1.** Comparison of Non3nExonFinder with selected existing tools relevant to CRISPR exon targeting.

| Feature | CHOPCHOP<br>(Labun et al.) | CRISPRware<br>(Malekos et al.) | KOnezumi<br>(Kuno et al.) | Non3nExonFinder |
| --- | --- | --- | --- | --- |
| Primary purpose | CRISPR gRNA design | Context-aware gRNA design | Mouse KO strategy design | Identification of shared non-triplet coding exons |
| Per-transcript exon/CDS annotation | △ | △ | X | O |
| Multiple transcripts analyzed jointly | O | O | △<sup>2</sup> | O |
| Exact shared exon-boundary detection | X | △<sup>1</sup> | X | O |
| CDS/UTR-aware exon classification | △ | △ | X | O |
| Transcript-specific coding contribution | X | X | X | O |
| Coding-length modulo-three evaluation | X | X | △<sup>2</sup> | O |
| Internal-exon filtering | X | X | X | O |
| Direct input of user-defined RefSeq transcript sets | △ | X | X | O |
| Interactive interface | O | X | O | O |

Note: O, explicitly supported; △, partially or indirectly supported; X, not explicitly described in the cited publication.
<sup>1</sup> CRISPRware can construct consensus gene models from exon and CDS features shared across isoforms, but does not explicitly require exact exon-boundary identity or quantify transcript-specific exon coding contributions.
<sup>2</sup> KOnezumi uses IMPC-defined critical exons that are common to all transcript variants and whose deletion is intended to disrupt more than 50% of the protein-coding sequence by frameshift, rather than performing de novo exon/CDS analysis of user-defined transcript sets.

**Fig. 1.**
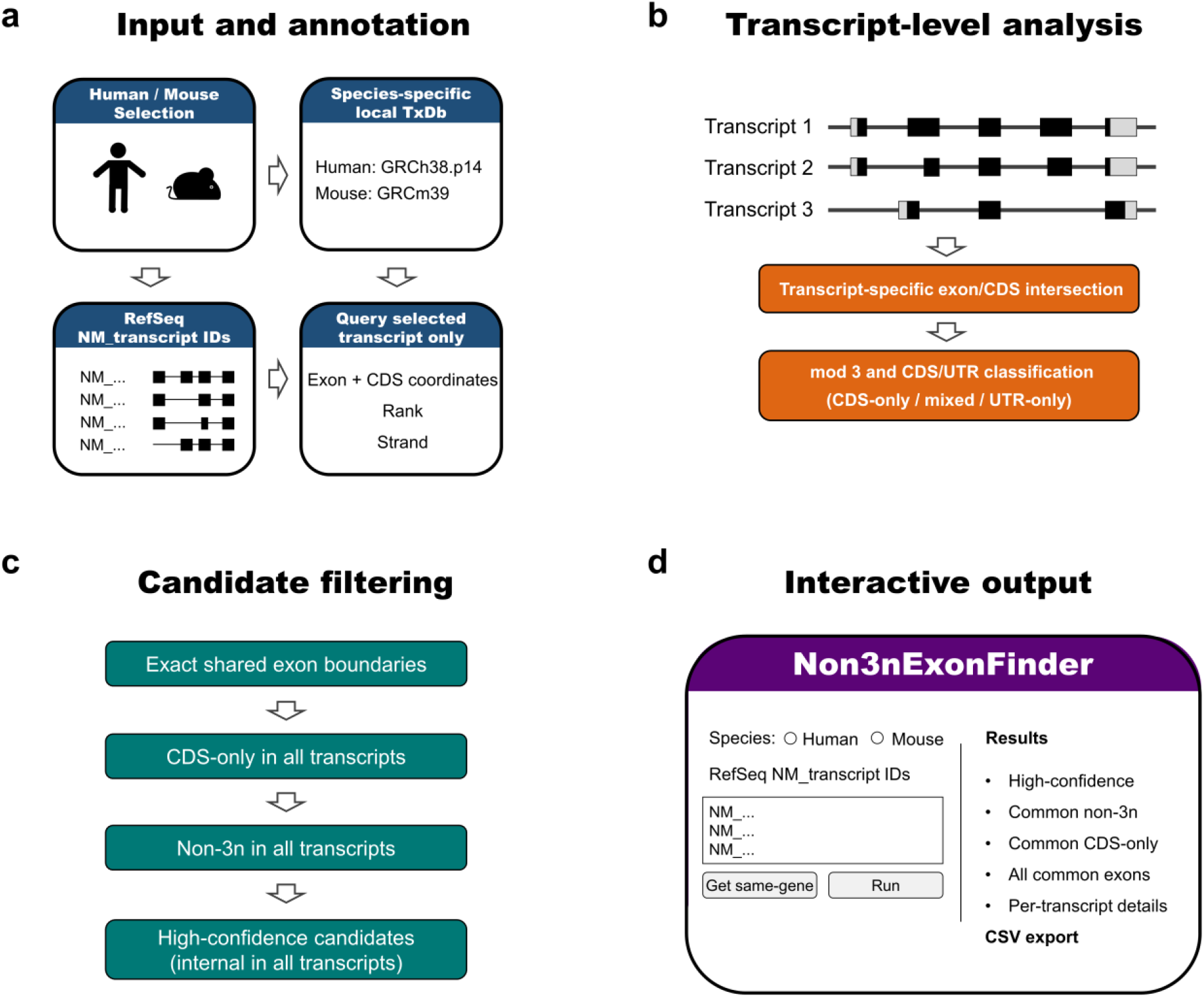
Overview of the Non3nExonFinder workflow. **(a)** Human or mouse RefSeq protein-coding transcripts are queried against species-specific local TxDb annotations. **(b)** Transcript-specific exon/CDS overlap is used to determine coding contribution and CDS/UTR classification. **(c)** Common exons are identified by exact genomic-boundary matching and subsequently filtered by CDS-only status, non-triplet coding contribution, and internal-exon status to identify high-confidence candidates. **(d)** The Shiny interface provides interactive result inspection and CSV export.

For each selected transcript, the program evaluates exon structure and coding contribution independently before integrating the results across the transcript set. Shared exons are progressively restricted to those that are CDS-only in all transcripts, have coding lengths not divisible by three in all transcripts, and are internal exons in every transcript. The resulting internal CDS-only non-triplet exons are reported as the most restrictive high-confidence candidate set (**Fig. 1b–d**).

### Application to a selected multi-transcript *TIA1* dataset

A representative analysis was performed using eight human RefSeq protein-coding transcripts of *TIA1*. All eight transcripts were retained in the GRCh38.p14-derived annotation, yielding 98 transcript-specific exon instances for analysis (**Fig. 2a**).

**Fig. 2.**
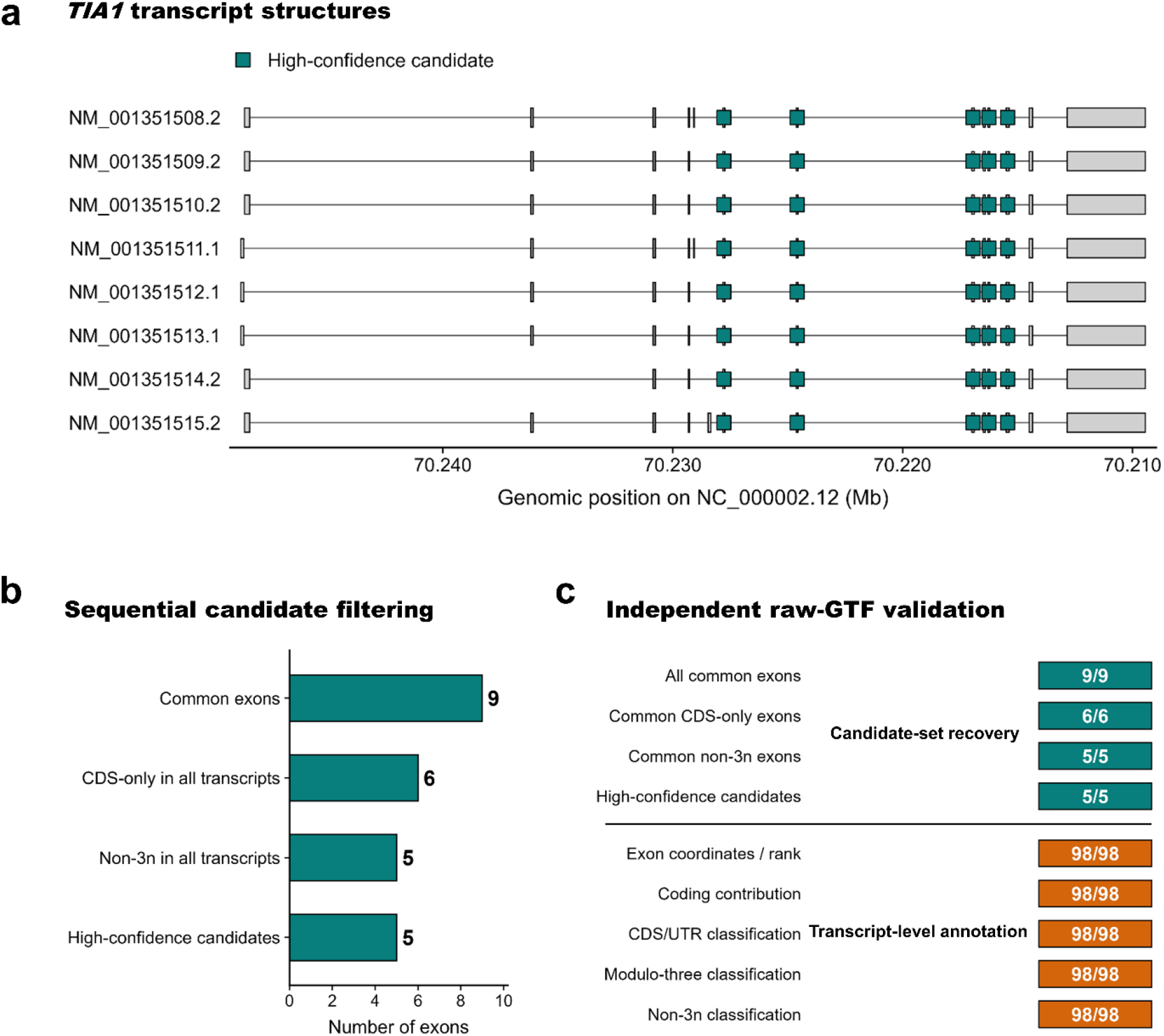
Application and independent validation of Non3nExonFinder using selected human *TIA1* transcripts. **(a)** Exon structures of eight RefSeq protein-coding *TIA1* transcripts, with high-confidence candidates indicated by teal markers. **(b)** Sequential filtering of shared exons identified nine common exons, six common CDS-only exons, five common non-triplet exons, and five high-confidence candidates. **(c)** Independent reconstruction from the original RefSeq GTF showed complete concordance for all 98 transcript-specific exon instances and exact recovery of all candidate sets.

Exact genomic-boundary matching identified nine exons shared across all eight transcripts. Six of these were CDS-only in every transcript, and five of the six CDS-only exons had coding lengths not divisible by three in all transcripts. All five non-triplet exons were also internal in every analyzed transcript and were therefore retained as high-confidence candidates (**Fig. 2b**).

Independent reconstruction from the original NCBI RefSeq GTF showed complete agreement with the Non3nExonFinder output. Exon coordinates and rank, transcript-specific coding contribution, CDS/UTR classification, coding-length modulo-three status, and non-triplet classification were concordant for all 98 transcript-specific exon instances. Independent candidate reconstruction also recovered the identical sets of nine common exons, six common CDS-only exons, five common non-triplet exons, and five high-confidence candidates (**Fig. 2c**).

### Computational performance

Computational performance was assessed using nested *CASP8* transcript sets containing 1, 5, 10, 20, and 40 RefSeq transcripts, with 10 timed replicate analyses performed for each transcript-set size. Elapsed time generally increased with transcript number, while analyses of the 40-transcript set remained below 10 s per run across the tested runs (**Fig. 3a**). These results indicate that transcript-specific exon/CDS analysis and sequential candidate filtering remained computationally tractable across the evaluated transcript-set sizes.

**Fig. 3.**
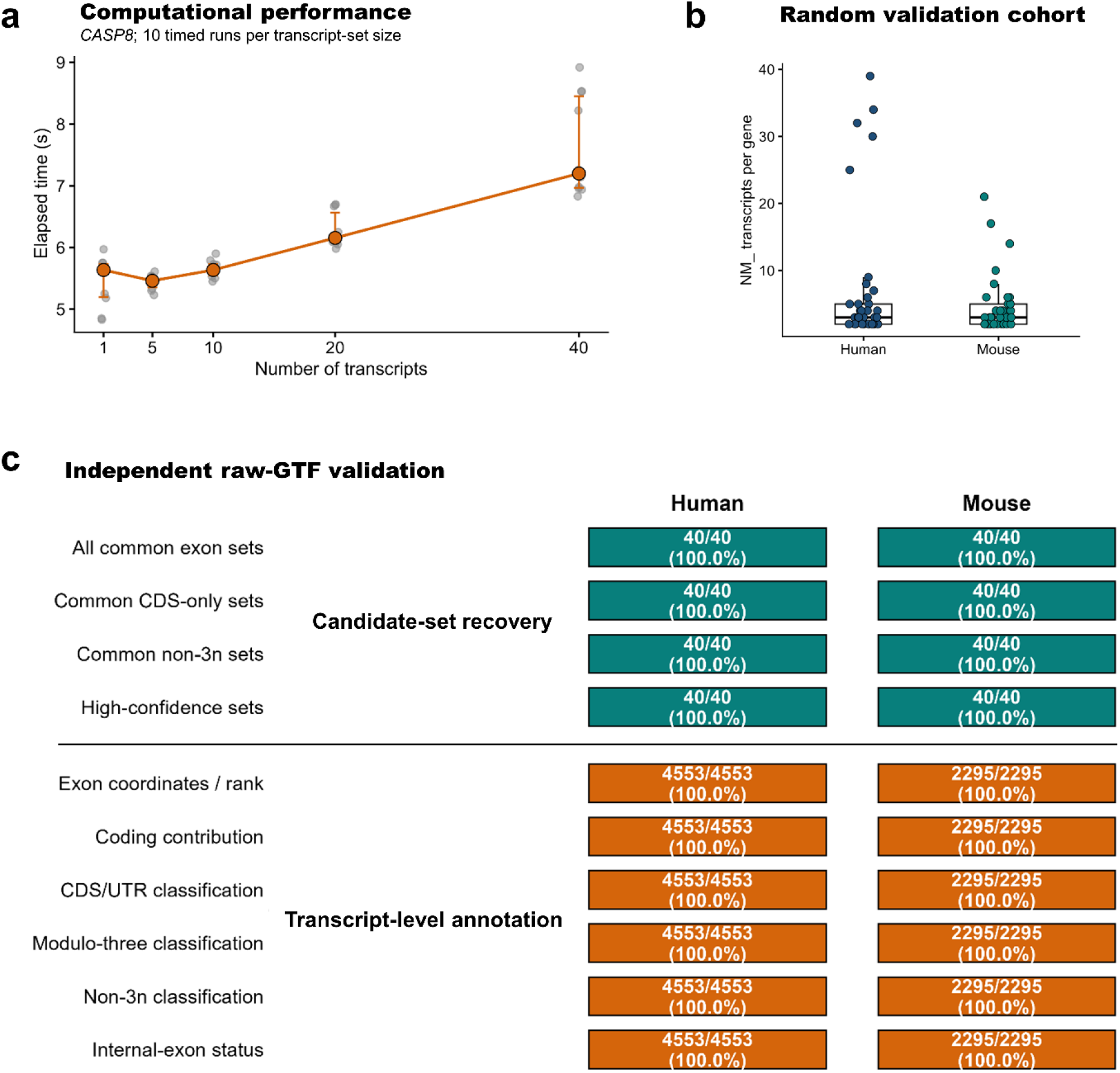
Computational performance and large-scale independent validation of Non3nExonFinder. **(a)** Runtime across nested *CASP8* transcript sets containing 1, 5, 10, 20, and 40 RefSeq transcripts. Points indicate individual runs, with median and interquartile range shown in orange. **(b)** Transcript counts of 40 randomly selected human and 40 mouse genes used for large-scale validation. **(c)** Independent raw-GTF validation showing exact recovery of all four candidate sets in all 40 genes per species and complete concordance for six transcript-level annotation features.

### Large-scale independent validation of Non3nExonFinder

Large-scale validation was performed using 40 randomly selected human genes and 40 randomly selected mouse genes, each containing at least two eligible RefSeq NM_ transcripts (**Fig. 3b**). The validation cohort comprised 276 human and 180 mouse transcripts, corresponding to 4,553 and 2,295 transcript-specific exon instances, respectively. In total, 456 transcripts and 6,848 exon instances were evaluated across the two species.

Transcript-level annotations were independently reconstructed directly from the original NCBI RefSeq GTF files. Complete concordance was observed for all six evaluated features: exon coordinates and rank, coding contribution, CDS/UTR classification, coding-length modulo-three status, non-triplet classification, and internal-exon status. All 4,553 human exon instances and all 2,295 mouse exon instances were concordant with the corresponding Non3nExonFinder outputs, with no exon-level discrepancies detected (**Fig. 3c**).

Candidate-set reconstruction produced the same result. For all 40 validation genes in each species, the independently reconstructed sets of all common exons, common CDS-only exons, common non-triplet exons, and high-confidence candidates were identical to the corresponding Non3nExonFinder outputs at every filtering stage (40/40, 100%). No candidate-set discrepancies were detected across the 80-gene validation cohort (**Fig. 3c**).

## DISCUSSION

Non3nExonFinder was developed to address transcript-annotation issues encountered when selecting coding exons for exon-deletion experiments. Users can retrieve additional RefSeq protein-coding transcripts associated with the same Gene ID as a specified transcript or analyze only a user-defined subset. Exon and CDS annotations are evaluated independently across transcripts, with shared exons identified using strict genomic-boundary criteria and assessed for their potential to disrupt the reading frame.

A key feature of the workflow is the distinction between genomic exon length and the actual coding contribution of an exon. Terminal or alternatively annotated exons may contain both coding and untranslated sequence, and applying a modulo-three calculation to the entire exon length could therefore misclassify the predicted coding consequence of exon removal. Non3nExonFinder addresses this issue by calculating exon-CDS overlap independently for each transcript and restricting its primary candidate set to exons classified as CDS-only across all selected transcripts.

The definition of a common exon is intentionally conservative. An exon is considered shared only when its chromosome, genomic start and end coordinates, and strand are identical across all selected transcripts. This criterion prevents partially overlapping or differently bounded exons from being treated as equivalent targets. Consequently, biologically related exon segments with transcript-dependent splice boundaries may be excluded, reflecting the intended purpose of the program: prioritizing a single genomic exon whose complete removal can be interpreted consistently across the selected transcript set.

Non3nExonFinder is designed as an upstream annotation utility rather than a complete CRISPR design platform and does not directly predict the functional consequences of exon deletion. Accordingly, a non-triplet coding exon should be interpreted as a candidate predicted to disrupt the downstream reading frame if the exon is successfully removed, rather than as a target guaranteed to produce a functional knockout. Because the program also reports coding-exon and CDS information beyond the non-triplet candidate set, these annotations may also be useful for evaluating target regions in other genome-editing strategies, including single-sgRNA knockout and knock-in approaches. For applications intended to achieve knockout through exon removal, the position of any resulting premature termination codon should additionally be evaluated in the context of transcript structure to assess the likelihood of nonsense-mediated mRNA decay (Kurosaki et al. 2019).

The current implementation has several limitations. Non3nExonFinder is restricted to human and mouse RefSeq protein-coding transcripts represented by NM_ accessions, and its results depend on the content and version of the TxDb annotation used for analysis. Results may therefore change as RefSeq transcript models are revised, and the biological relevance of the transcript subset included in an analysis remains dependent on user selection.

## CONCLUSION

Non3nExonFinder is a transcript-aware upstream annotation tool for identifying shared non-triplet coding exons across human and mouse RefSeq transcripts. Independent validation showed complete concordance across 6,848 transcript-specific exon instances and candidate sets from 80 genes. These results support the use of Non3nExonFinder as a reproducible annotation framework for prioritizing exon-deletion targets before downstream CRISPR guide design and experimental validation.

## DATA AVAILABILITY

The source code for Non3nExonFinder and all data underlying the validation and computational performance analyses are publicly available on GitHub (https://github.com/sungyeonlee0711/Non3nExonFinder) and Zenodo (https://doi.org/10.5281/zenodo.22884913). The repository includes the *TIA1* validation dataset, large-scale validation gene and transcript sets, exon-level validation results, candidate-set comparisons, gene-level validation summaries, computational performance benchmark data, quality-control outputs, and session information. The interactive web application is available at https://sungyeonlee0711.shinyapps.io/Non3nExonFinder. The original human and mouse RefSeq annotation files used in this study are publicly available from NCBI under the accessions GCF_000001405.40 and GCF_000001635.27, respectively.

## AUTHOR CONTRIBUTIONS

**Sung-Yeon Lee:** Conceptualization, Writing – review & editing, Writing – original draft, Project administration, Visualization, Validation, Resources, Supervision, Software, Methodology, Investigation, Formal analysis, Data curation. **Joo-Hee Lee:** Investigation, Validation.

